# Low polyethylene glycol (PEG) concentration contributes to photoprotection in *Chlamydomonas reinhardtii* under highlight stress

**DOI:** 10.1101/2022.08.30.505872

**Authors:** Jerome Xavier, Ranay Mohan Yadav, Rajagopal Subramanyam

## Abstract

The highlight is one of the major problems encountered by all the autotrophs in the natural environment, as it affects the photosynthetic performance of these organisms, they were equipped themself with the diverse photoprotective mechanism. However, the drastic climate change in recent years had a more lethal effect on these organisms as it is beyond their threshold to withstand. In the past decade, scientists have unravelled many photoprotective mechanisms like qE, qZ, qT and qI (NPQ) used by autotrophs to combat highlight stress and the thirst for discovering such a new mechanism remains constant. In this regard, we studied the effect of mild osmotic stress (2%PEG) in alleviating high-light stress using *Chlamydomonas* (C) *reinhardtii* as a model system. The cells were grown in low Polyethylene glycol (PEG)-induced osmotic stress at varying light intensities; their response to these treatments had been examined via biochemical and biophysical approaches. The PEG-treated cells showed better growth and photosynthetic efficiency even at highlight than control, but their NPQ level is lesser than control, suggesting a unique or novel photoprotective mechanism is operating in PEG-treated samples. The Circular dichroism and ATG8 localization assay suggest that supercomplexes organization is not much disturbed in PEG-treated samples irrespective of light intensity. Also, the latter indicates that PEG-treated cells were healthier than the control at the highlight. This result is promising as we can improve the algal biomass under natural environmental conditions with fluctuating light intensity. As algal biomass has immense commercial importance in biofuel production, cosmetics and pharmaceutical application. This mechanism can be exploited to promote the socio-economic status of our nation.

**Highlights:**

1. PEG application triggers ROS accumulations, stimulating signalling cascade and overcoming highlight-induced compromise in biomass.
2. Under highlight, the photosynthetic pigment (chlorophyll and carotenoid) constantly increased with PEG except for C-250. Specifically, the photoprotective pigment like violaxanthin and lutein has been increased.
3. PEG shielding stabilizes pigment-protein interaction, confirming that the supercomplex organization is in its native state.
4. PEG keeps low ATG8 accumulation, preventing cells and enabling it to combat highlights more effectively.

**Graphical abstract:** 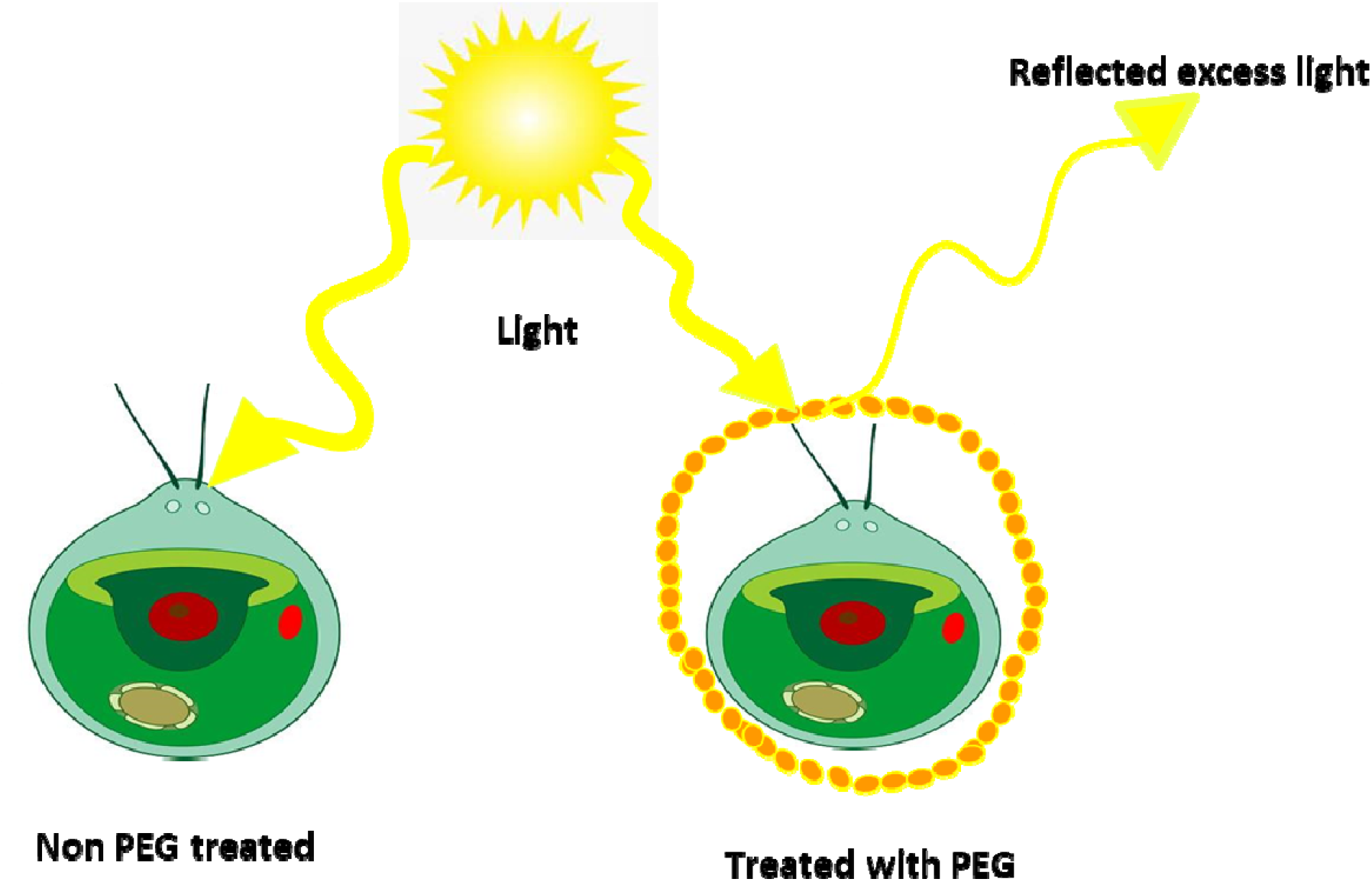

## Introduction

All life on Earth comprises a basic carbon skeleton and inorganic elements like oxygen, hydrogen, etc. Even crucial biomolecules like carbohydrates, protein, and nucleic acids are derivatives of atmospheric CO_2_ and H_2_O. Photosynthesis is the only process that can fix atmospheric carbon dioxide as a carbohydrate and supply it to the entire biosphere. Hence it is the vital process that drives the energy cycle to support life (Madireddi et al., 2014). However, the photosynthetic yield will always tend to fluctuate due to environmental conditions caused by biotic and abiotic factors. Among the abiotic factors, drought and highlight affect photosynthetic performance to a maximum since optimum light and water are the two main requirements for effective photosynthesis. The efficient conversion of light to biomass is photosynthetic efficiency(Vejrazka et al., 2011). In natural selection, only organisms that adapt to the available resources survive.Photoprotection is one such adaptation against extreme highlights (Erik H Murchie et al., n.d.). In a natural environment, photosynthetic organisms are not always exposed to optimum light intensity or fluctuation. High light leads to over-reduction of the photosystem resulting in superoxide generation, which causes photoinhibition. Reduction in LHCS, disruption to the photosynthetic super complex organization, and degradation of reaction center protein subunits are the most common changes in *Chlamydomonas reinhardtii* while exposed to high light(Nama et al., 2015). Hence light absorption is tightly regulated by various photo-acclimation processes(Ware et al., 2015) like NPQ, state transition, and alteration of photosynthetic protein turnover(Neelam et al., 2013).

Apart from these mechanisms, two proteins, namely Psbs and LHCSR3, were over-expressed only during high light exposure, helping regulate energy balance and dissipation. The potential behaviour of autotrophs to acclimatize with their growth environment amidst multiple stress factors with the expense of least energy has always amazed the scientific community. As photoprotection and photochemistry share inverse relations, scientists thought of manipulating photoprotection to improve photosynthetic yield (E H Murchie et al., 2009). Polyethylene glycol is a non-ionic water-soluble polymer that can mimic drought stress by reducing water potential, thereby creating osmotic stress to organisms under study. Its chemically inert behaviour makes it ideal for inducing osmotic pressure in biochemical experiments(Ahmad et al., 2020). Our previous observation shows that among various concentrations of PEG treatment, 2% promoted algal growth compared to higher concentrations. Hence in this study, we are interested in finding whether the PEG molecule offers any photoprotection. If so, what kind of photoprotection is it? We grew Chlamydomonas reinhardtii in mild PEG(2%) induced osmotic stress along with different high light intensities. Their response to these treatments has been examined via biochemical and biophysical approaches.

## Results

### Growth vs ROS in light and PEG-treated cells

The photosynthetic organisms, under environmental stress primarily generates reactive oxygen species (ROS) (Nagy et al., 2018). Reactive oxygen species are essential signalling molecules of any cell. Under mild stress conditions, photosynthetic organisms regulate ROS production by increasing ROS scavenging activity (Laloi et al., 2004), induction of NPQ (Erickson et al., 2015), and upregulation of alternative electron transport pathways (Erickson et al., 2015). Under 2% PEG, the growth of *C. reinhardtii* cells was similar to that of control cells but Increased PEG concentrations have reduced growth and cell size. It is evident from this observation that osmotic stress affects the growth and morphology of *C. reinhardtii* at higher concentrations of PEG. Therefore we chose to work with cells grown with 2% PEG under different light conditions. The growth curve analysis shows there is not much difference in cell density till 24 hrs hence considered as lag phase. The difference in growth pattern occurs after 24 hrs, as represented in Figure 1a. There is a decline in the growth momentum with an increase in light intensity in both conditions, but PEG treated sample’s growth rate is better than the non-treated at respective light intensities. Among them, 250 µmol m^−2^ s^−1^ light intensity is very favourable for cell growth in both PEG treated as well as non-treated. It is interesting to find that the 1000 µmol m^−2^s^−1^ PEG treated sample remains in LOG while all other cultures attain a stationary phase. The 2′-7′-Dichlorodihydrofluorescein diacetate (H2-DCFDA) is a cell-permeable fluorescent dye used to quantify intracellular ROS levels. Within the cell, H2-DCFDA gets converted to H2-DCF by esterases, which will be oxidized by ROS, thereby giving a fluorescence signal detected by the confocal system (Eruslanov et al., 2010). The Confocal microscopic image shows that ROS generation increases in high light intensity in both PEG treated as well as in light samples. However, their accumulation is comparatively higher in all PEG samples than in the control. Earlier reports suggest ROS participate in retrograde signalling for inducing high light-responsive genes in *Nicotiana benthamiana* (Exposito-Rodriguez et al., 2017). Hence, more elevated ROS may support PEG samples to combat highlight stress.

**Figure 1:**
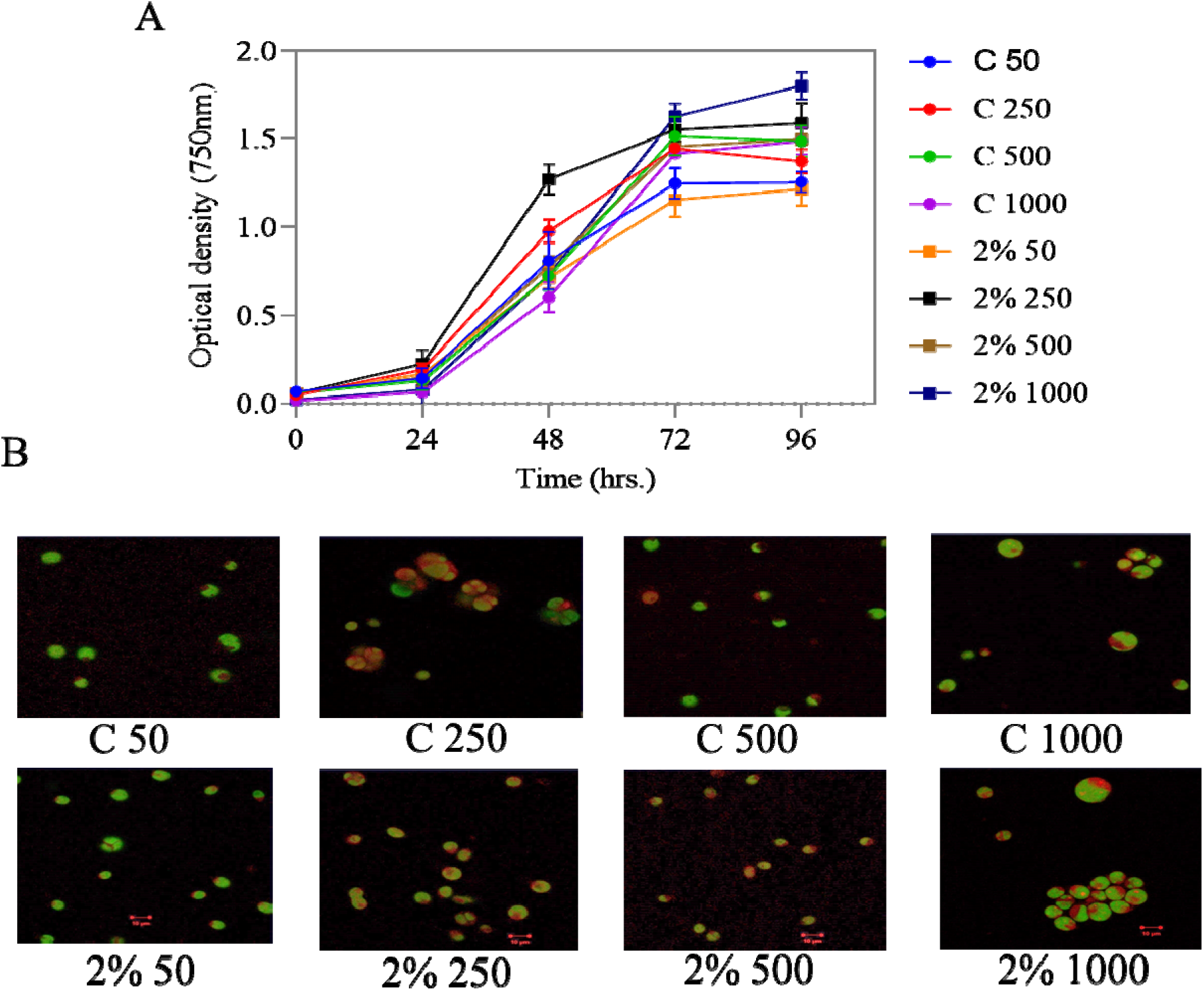
Growth physiology of all culture conditions under study, (A) cell growth measured at regular time interval using PerkinElmer UV visible spectrometer at 750 nm, (B) qualitative estimation of Reactive oxygen species using H_2_DCFDA dye in Carl Zeiss NL0 710 confocal microscope.

#### Changes in Chl and carotenoid pigment content in light and PEG-treated sample

Pigment estimation for each sample was done and plotted in the graph (**Figure 2**). We found that total chlorophyll concentration (**Figure 2A**) in control samples increases with increasing light intensity C-250 is the highest. Whereas in PEG treated samples, chlorophyll concentrations were higher and almost similar in all three samples 2%PEG-250, 500 and 1000. The slight decrease in 2%PEG-250 chlorophyll concentration to its control is due to low chlorophyll-a concentration. The chlorophyll a/b ratio (**Figure 2B**) shows the correlation between C-50 and C-250, whereas, in PEG samples, there are no significant changes. The increase or decrease in these ratios is mainly contributed by chl-a and b concentrations. The total carotenoid concentration data was shown in (**Figure 2C**); here C-250 sample possesses a very high carotenoid concentration among control samples; in PEG samples, a gradual increase of carotenoid is observed in highlight still it is not up to the mark of control samples. To quantify and identify the presence of carotenoids expressed under light and PEG stress, we have run the samples in HPLC. All samples contained Chl *a*, Chl *b*, β-carotene (β - Car), violaxanthin (Vio), lutein (Lut), zeaxanthin (Zea), and xanthophyll (Xan) at a different stage of HPLC run, although their relative abundance varied under high light with PEG.

**Figure 2:**
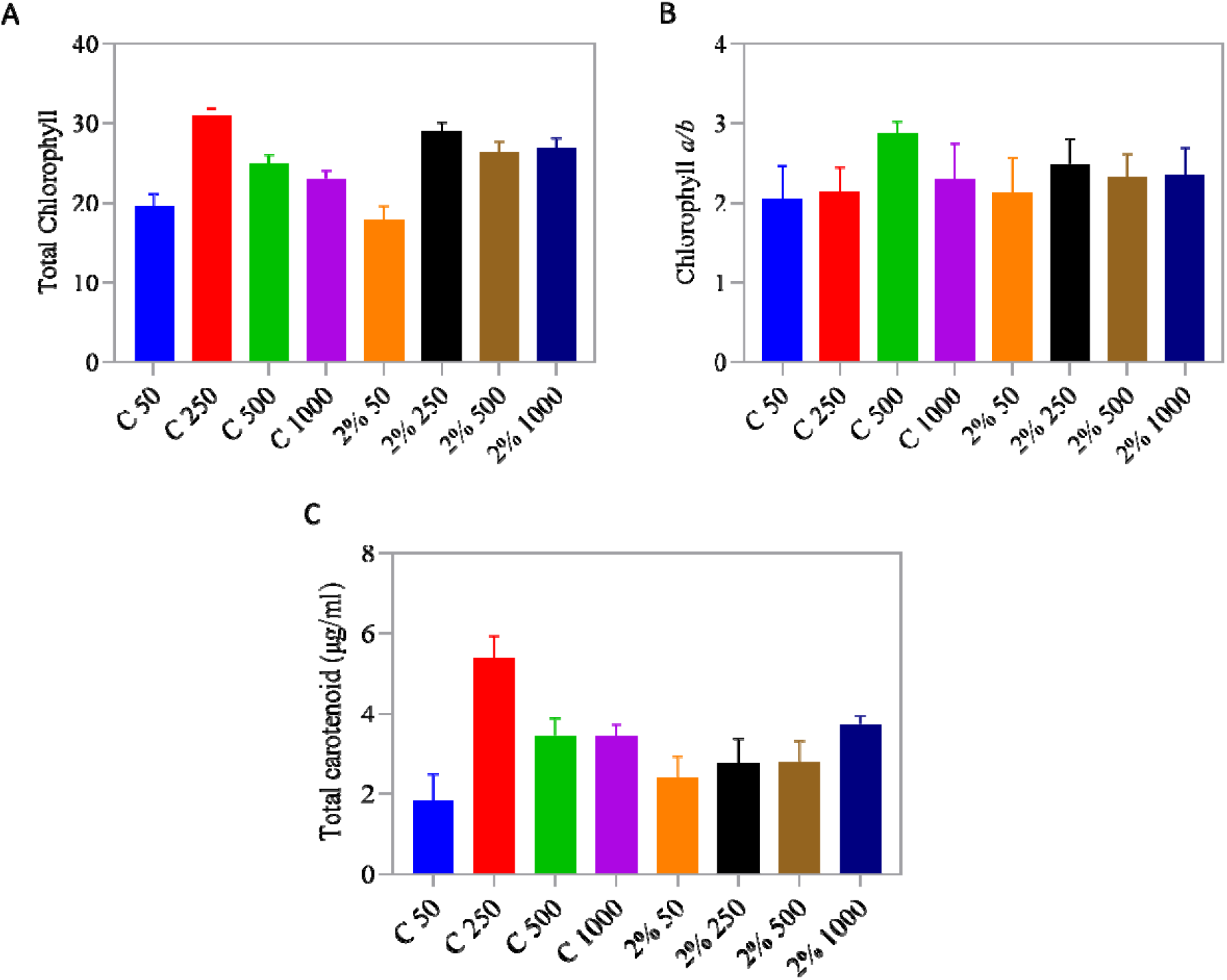
Photosynthetic pigment estimation using PerkinElmer UV visible spectrophotometer and concentration is calculated by adopting equations of lichtenthaler(1987) and Porra et al. (1989). (A) graphical presentation of total chlorophyll contents,(B) shows the variation in Chl a to Chlb in all culture conditions, (C) Carotenoid concentration has been reported.

#### Effect of light and PEG stress on Chl a fluorescence and photochemical activity

Fast Chlorophyll *a* fluorescence transient OJIP analysis, reflects the reduction of the electron transport chain; hence, it is used to describe photosynthetic yield performance. The chlorophyll fluorescence reflects the photochemical activity of PS II and the redox status of plastoquinone. Each stage in this transient reflects a particular phase in the electron transport system: The O-J phase in the transient is linked to QA reduction (Schreiber et al., 1987), and the rise from the J-I phase is due to the accumulation of QA- to QB- and subsequent QA - to QB and O to P raise in the fluorescence yield reflecting the variable fluorescence raise Fv is due to reduction of PQ pool size (Khan et al., 2021). The typical fluorescence transient curve of wild-type culture C-50 was shown in (**Figure 3A**); comparing it with the rest showed variation in their transient curve. The cultures C-250, C-500, 2%PEG-50, 2%PEG-250 and 2%PEG-500 deviate from C-50 at the J-P phase, whereas cultures C-1000 and 2%PEG-1000 deviate in the entire O-P phase. The photochemical yield Fv/Fm ratio (**Figure 3B**) had decreased among control samples grown at high light, whereas in the case of PEG treated samples except 2%PEG-1000, the photochemical yield remains stable almost equal to the C-50 sample.

**Figure 3:**
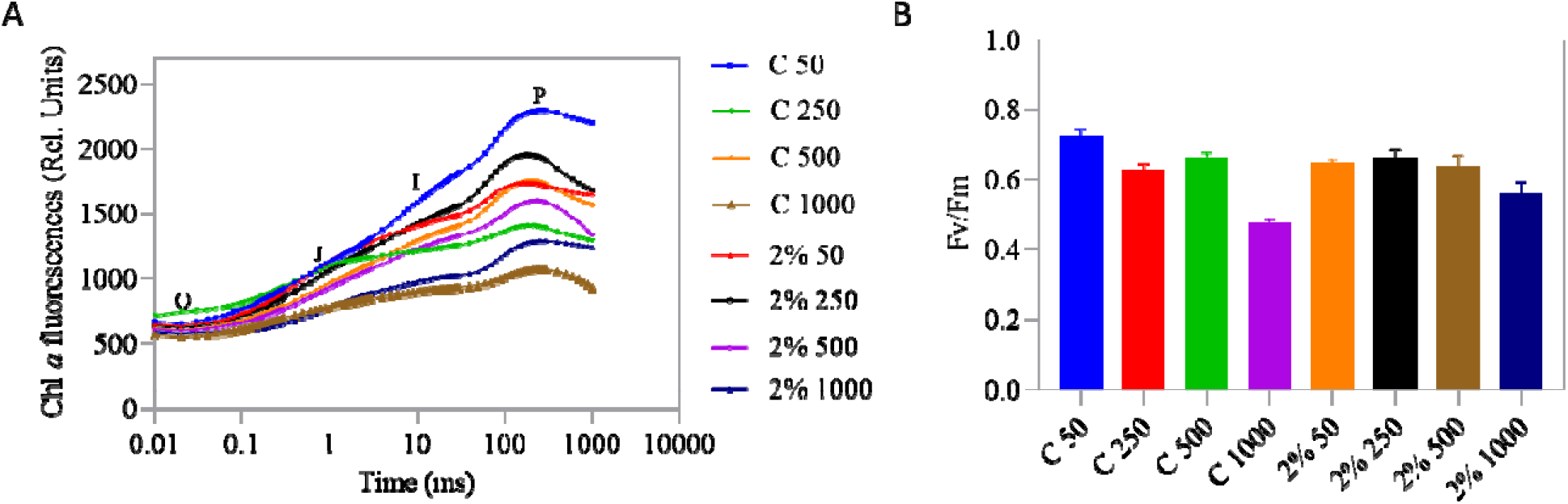
shows the typical OJIP fluorescence transients parameters(A) and photochemical yield Fv/Fm (B) were calculated by Plant Efficiency Analyzer (Hansatech Instr. Ltd, Kings Lynn, Norfolk, UK) for one second with excitation light wavelength at 650nm in liquid cell cultures.

#### Change in pigment-protein thylakoid complexes in PEG-treated sample

Circular dichroism is a biophysical technique to determine the structural change of pigment– pigment interactions and macro-organization of supercomplexes. It indicates the difference in absorption of left-handed and right-handed circularly polarized light; hence any pigment-protein interaction shift can be detected by analysing it. CD spectroscopy of light and PEG-treated thylakoid membranes showed the peaks in the visible region (400 - 800 nm) and has been divided into the psi-type band (650 - 695 nm) and soret region (440 - 510 nm) (**Akhtar et al**., **2015**). The absorption spectra depict the denaturation or degradation of pigments and supercomplexes; almost every control sample’s spectrum is uniform. In contrast, in PEG-treated samples, deviation occurs in every sample. CD spectra of all the cultures were shown in (**Figures 4A and B**); while analysing all four control samples(C-50, C-250, C-500 and C-1000), there is a remarkable decline in chl a related prominent positive peak along with slight shifting towards the soret region side occurred at 680nm as the light intensity increases. The opposing band at 672nm also got disturbed in high light-exposed samples, but the chl b related spectrum (640 and 460nm) remains consistent in all the control samples, irrespective of different light intensities. Whereas among PEG treated samples, even though we can observe significant changes in chl *a* and *b* peak, it is not as drastic as controls. The CD data suggest pigment-protein interaction i.e. chlorophyll-thylakoid membrane is more stable in PEG samples.

**Figure 4:**
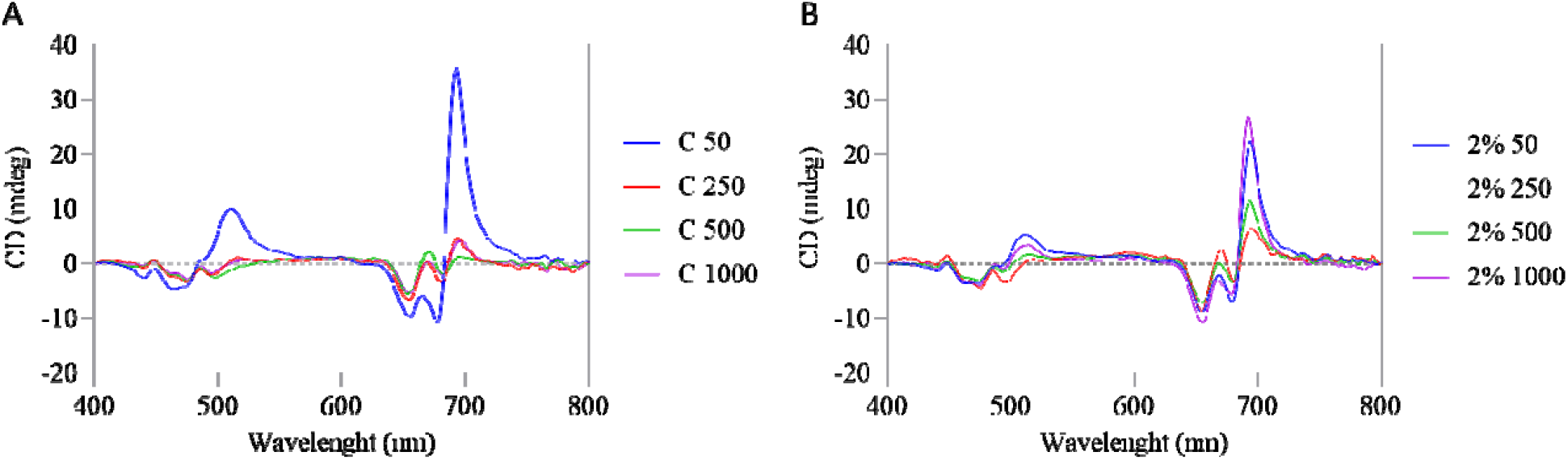
Absorption and CD spectrum of cell suspension culture using circular dichroism spectrophotometer. The peaks at 443(+) nm, 680 nm(+), and 672(-) nm are from chl *a*, wherea peak at 640nm(-) and 460nm(-) is from chl b. The positive band around 512 and 418nm is from the carotenoid.

#### Relationship between time-based NPQ measurement and LHCSR3 accumulation in light and PEG cells

The Non-photochemical quenching (NPQ) helps dissipate excess light energy as heat, which is triggered by sensing the pH variation of the lumen. Logically, high light treated samples should have more NPQ since their lumen acidification is more. All control samples show higher NPQ levels than PEG-treated samples with C-50 and its 2%PEG-50 as an exception (**Figure 5 A, B**). The LHCSR3 is a homologue of PSBS protein of plants having a role in energy-dependent quenching qE a component of NPQ (Xuey et al., 2015). The immunoblot analysis for LHCSR3 showed no expression in low light because of low lumen acidification, which were in agreement with the NPQ data. Further, their expression is lesser in PEG samples than in controls indicating lesser exposure to stress in PEG samples. Here, the PsaF protein is used as the loading control.

**Figure 5:**
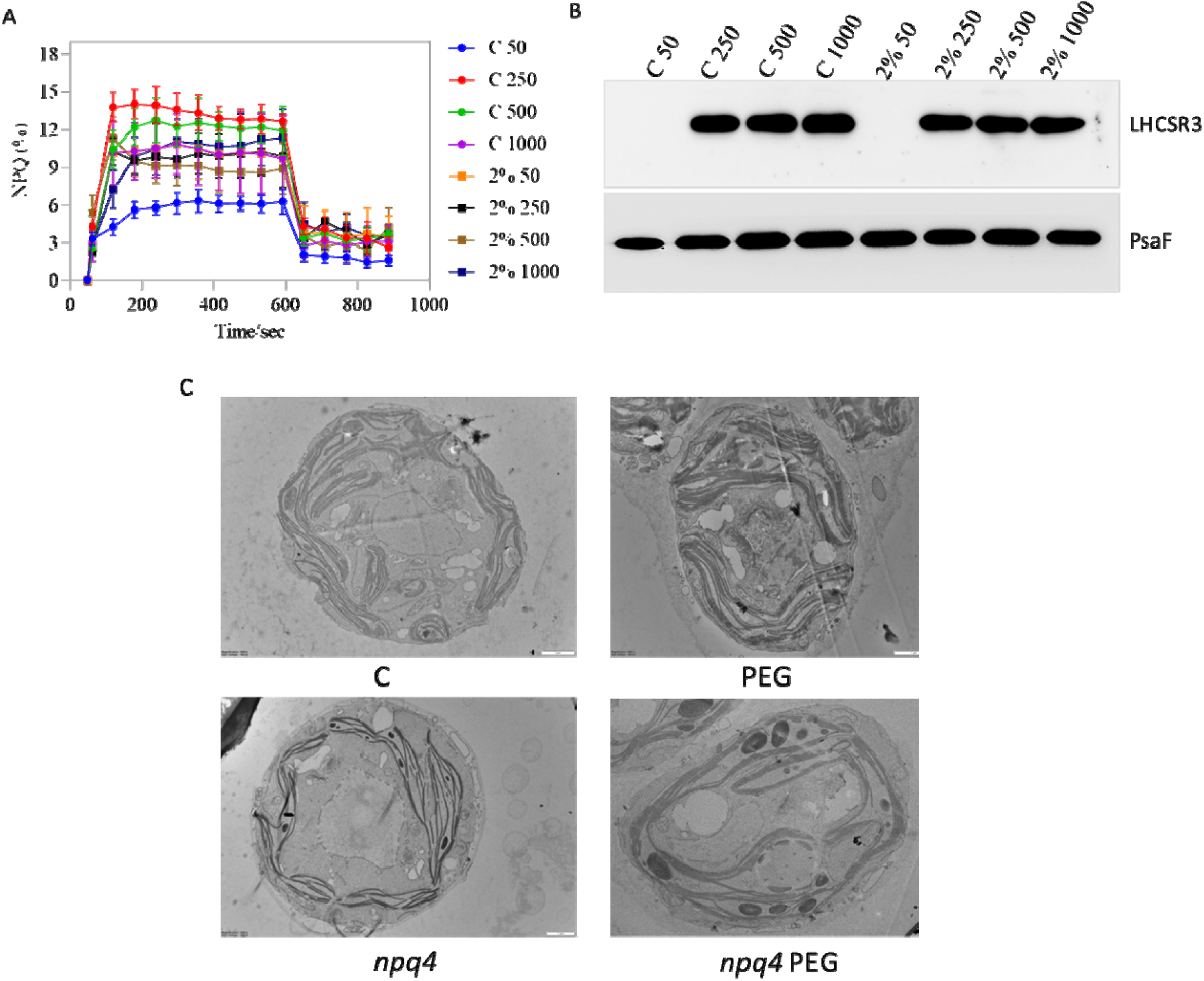
Represents various biophysical data related to photosystem II like: (A) Non-photochemical quenching, (B) immunoblot showing LHCSR3 expression and (C) TEM imag analysis showing the cell morphology.

#### Light and PEG change the PSII photosynthetic parameters

On continuation of NPQ, various other biophysical parameters have been investigated; The electron transport rate (**Figure 6A**) around PS II shows all the control samples except C-1000 have higher PS II yields than their respective PEG-treated samples. PS II yield also gave the same result as ETRII (**Figure 6B**). In the case of non-regulated heat dissipation, the results are vice versa to yield and ETR II, and here the PEG-treated samples show higher readings than controls (**Figure 6C**). The NPQ yield suggests variation from the previous pattern of (YII); here, C-50 and C-500 seem to be greater than the PEG samples (2%PEG-50 and 2%PEG-500) (**Figure 6D**).

**Figure 6:**
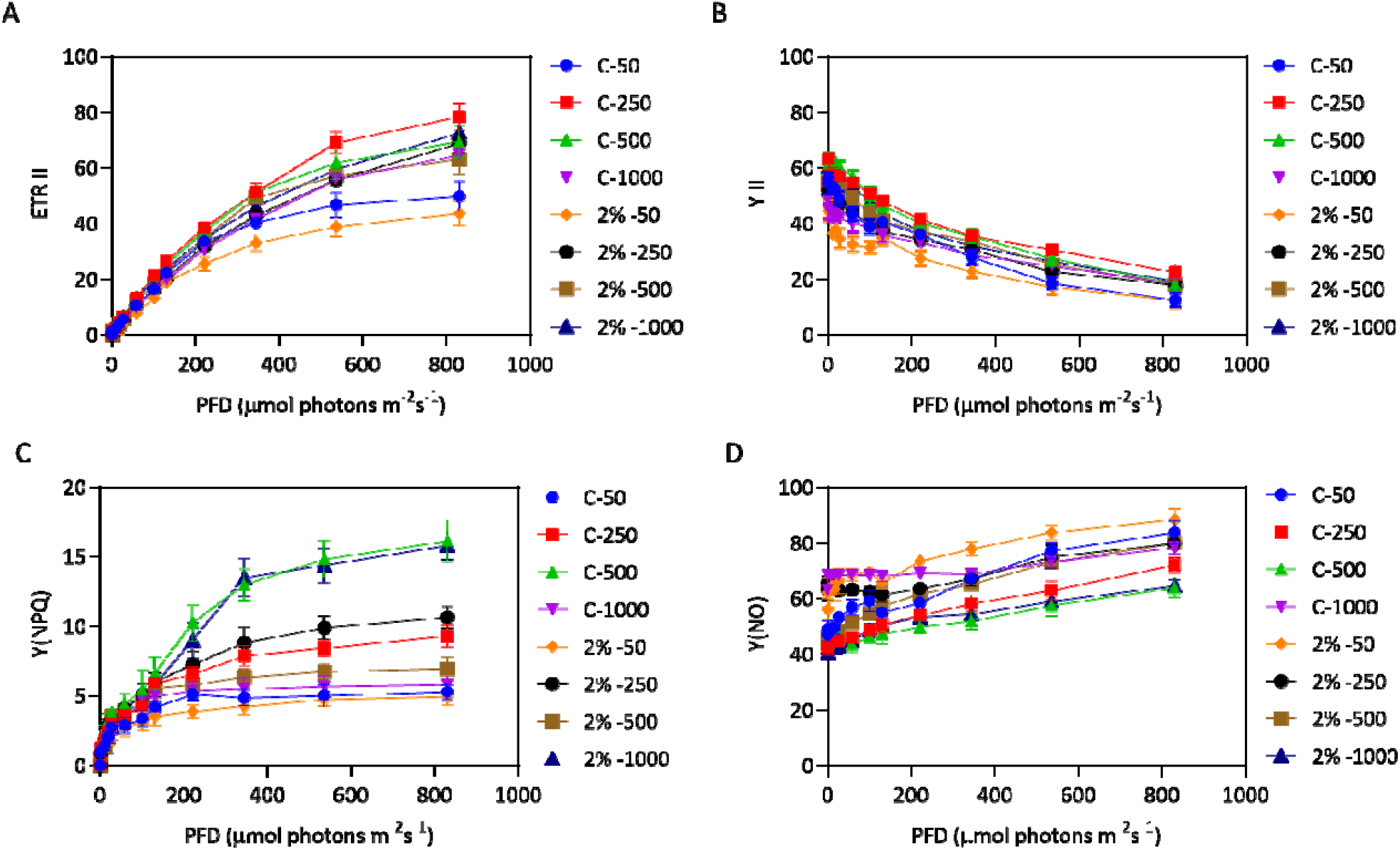
Represents various biophysical data related to photosystem II like: (A) Electron transport rate across PS II, ETRII (B) Photosystem II yield of individual culture conditions,YII (C) Yield of non-photochemical quenching. (D) Yield of non-regulated energy dissipation.

**Figure 7:**
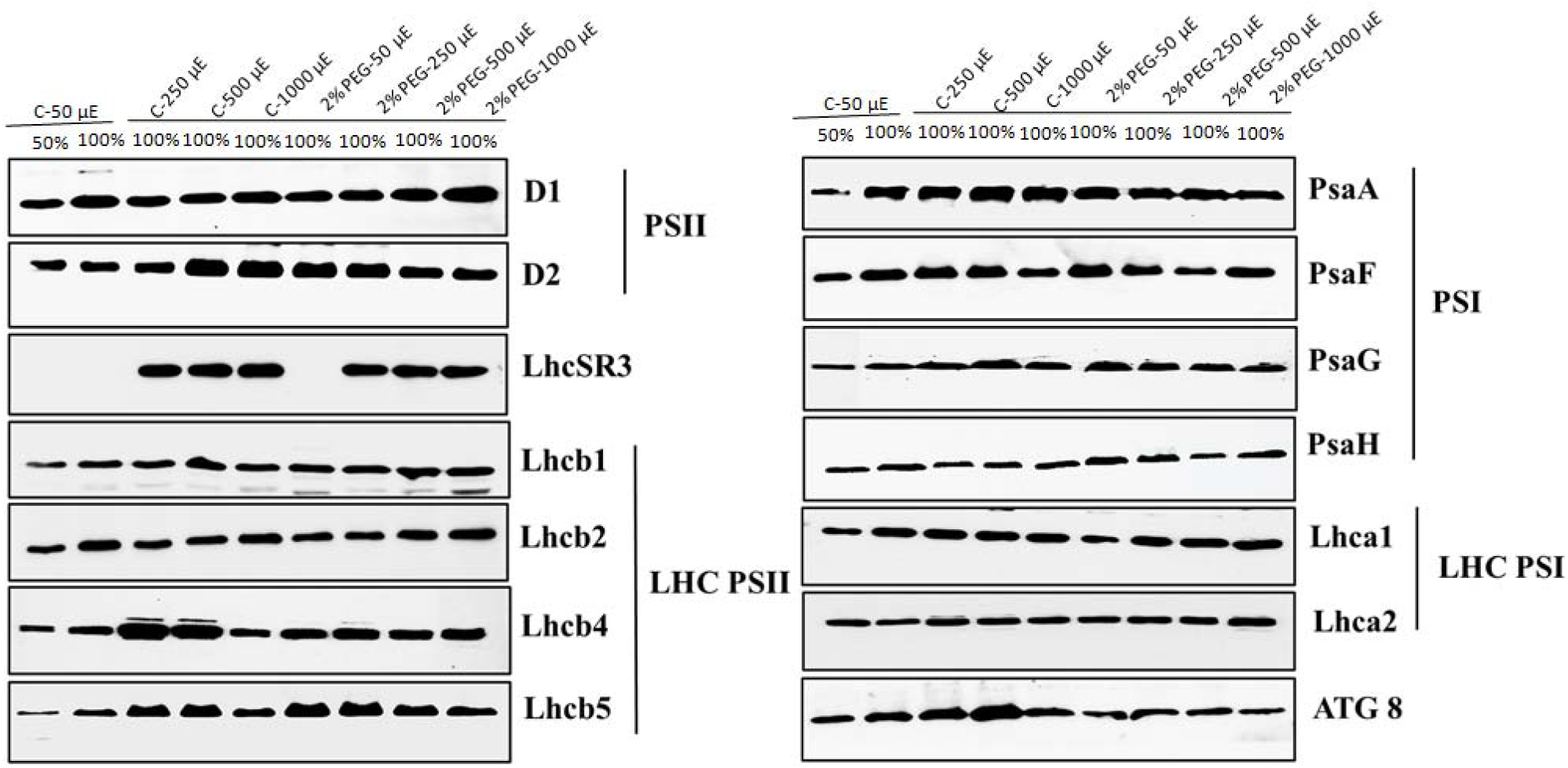
*C*.*reinhardtii* cultures under different conditions, proteins were separated based on their molecular weight by SDS-PAGE by taking equal chlorophyll concentration from each sample to normalize, followed by Immunoblot analysis of their thylakoid membrane proteins.

#### Immunoblot analysis of photosynthetic proteins

We have performed the immunoblot analysis of PS core proteins, D1, D2, PsaA, PsaF, PsaG and PsaH, along with major antenna proteins CP43 and CP47 and the PSII and PSI light-harvesting complex proteins Lhcb1, Lhcb2, Lhcb4, Lhcb5, Lhca1 and Lhca2. The immunoblot analysis showed four LHCs of PS II, Lhcb1 and Lhcb2; protein content was stabilized in all culture conditions, but in Lhcb5, the accumulation was higher in C-250, C-500, 2%PEG-50, 2%PEG-250 and 2%PEG-500 than even its normal growth conditions C-50. Further expression was comparatively higher in C-250 and C-500 in the case of Lhcb4. There is slight protein content variation among the two PS I LHCs (Lhca1 and Lhca2). We analysed two stress-related protein expression levels in all these conditions; the LhcSR3 expression occurred only in highlight conditions (250, 500 and 1000μmol m-2 s -1) in both control and PEG treated samples. The apoptosis-inducing protein ATG8 expression was higher in all control samples than in PEG-treated samples.

#### Subcellular Localization and ATG8 protein accumulation in light and PEG-treated cells

The process of autophagy always reflects the health status of a cell; even though there are more than 40 ATGs involved in the entire autophagy process, scientists prefer to use ATG8 as the marker to asses the cell death rate (Jacquet et al., 2021) The sub-cellular localization of ATG8 protein was done using an anti-ATG8 antibody and visualized in a confocal immunofluorescence microscope (**Figure 8A**). Atg8 expression was seen in both PEG as well as control samples, but intense observation shows that all PEG-treated samples had less ATG8 expression than their controls even at the highlight; the immunoblot analysis (**Figure 8B**) using anti-ATG8 antibody showed a similar result.

**Figure 8:**
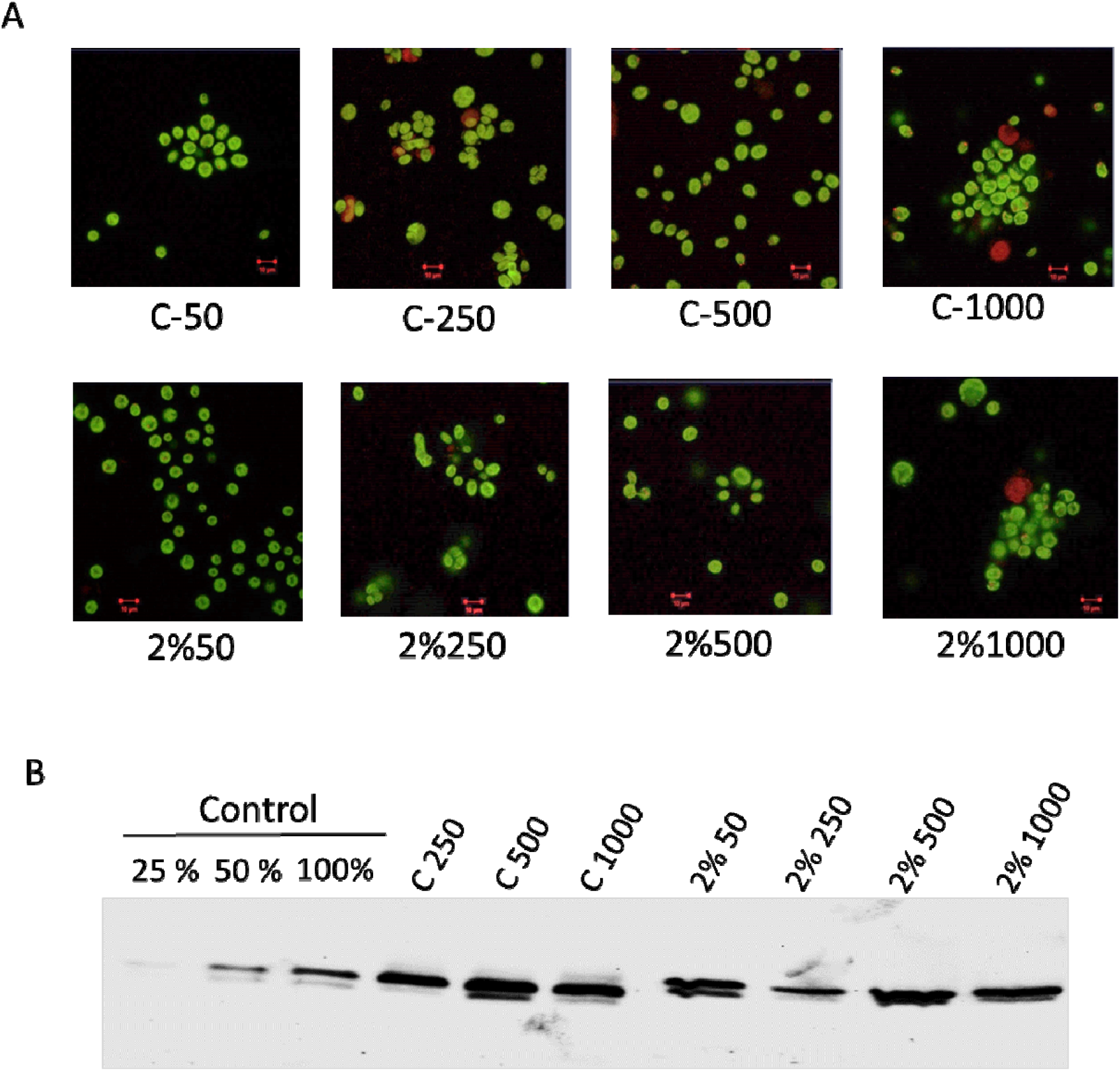
Shows ATG8 expression in all the sample conditions, Cells were collected after 3^rd^ day and immunoassay with anti-ATG8 antibody. (A) Chloroplasts were visualized by auto-fluorescence (in green). The image was taken using a Zeiss confocal microscope.(B)Immunoblot analysis for ATG8 to support the confocal localization data.

## Discussion

Stress, irrespective of its nature and intensity, always affects autotrophs; hence, most developed mechanisms to counteract and overcome such adverse conditions (Zhang et al., 2022). In this study, we report C. reinhardtii cell growth, photochemistry and super complex organisation under the combination of low osmotic stress and various light intensities (50, 250, 500 and 1000 μmol photons m^−2^s^−1^). Water and light are two major factors essential for autotrophs as they affect photosynthesis. We expected a severe effect on the growth of cells subjected to both of these stresses (osmotic and highlight stress). The previous report suggests that PEG-induced osmotic stress compromised the biomass in Sorghum (O’Donnell et al., 2013). Surprisingly we found no such compromise in growth even at highlight; further growth is better in treatment than their respective controls without PEG. It seems the mild osmotic stress is helping the cells to alleviate the highlight stress. The accumulation of high ROS levels in PEG-treated cells compared with control cells might act as signal transduce activating signalling cascade like mitogen-activated protein kinases (MAPKs) to combat highlight stress (Nadarajah, 2020). The carotenoid pigment has a crucial role as an antioxidant under stress conditions; mutant of carotenoid biosynthesis showed bleached phenotype(Nisar et al., 2015). The pigment estimation, particularly carotenoid content, was significantly reduced in PEG-treated cells was unexpected as it ruled out the possibility of a carotenoid role in photoprotection which usually occurs at the highlight.

The photochemical yield (F_v_ /F_m_) shows that PEG-treated cells were photosynthetically more active than their respective controls (Fig.3c). The fast chlorophyll *a* fluorescence transient reflects the successive reduction of the PQ pool of PSII. The OJIP curve for all the samples was plotted (Fig.3d), except for the control cells exposed to low light intensity (50 μmol photons m^−2^s^−1^); others exhibited abolition of IP phase due to reduction in transfer of an electron from PSII to PSI. However, the level of compromise is lesser in PEG-treated samples compared with their controls supporting the F_v_/F_m_ ratio data. Since significant photosynthetic pigments like chlorophyll and carotenoid were found in chloroplast’s thylakoid, evaluating their interaction will help assess the cell’s physiological status. CD spectrum is a sensitive technique that records any changes in the structure or arrangement of these pigment-protein complexes (Garab et al., 2009).

While analysing the CD spectrum of the isolated thylakoid membrane of *C. reinhardtii*, both PEG treated and controls each subjected to different intensities of highlight (Fig .4). We found that both Chlorophyll a and b corresponding peaks got much disturbed at high light intensities only in case-control samples. Among them, chl a peak disturbed to a greater extent which might be due to disruption of chl *a* in PSI-LHCI super complex due to high light. In the case of PEG-treated samples, we can observe this disruption of pigment-protein interaction, but their level is significantly low compared to their respective controls at the highlight. The increase in NPQ upon increasing light intensity is observed in both control and treatment, which is logical as it is a photoprotective mechanism involved in dissipating excess energy. However, the NPQ rate is low for all PEG treated samples than for controls which are supported by the higher ROS accumulation in PEG samples (Fig.1). This shows that some novel photoprotection mechanism operates other than NPQ, particularly in all PEG samples which protect them from high light-induced photodamage. The higher Non-regulated energy dissipation in PEG samples adds strength to our assumption. The immunoblot analysis of PSI and II-related proteins shows not much variation between control and treated samples. However, ATG8, an apoptosis-inducing protein expression, was much higher in control cells than PEG treated, suggesting the PEG treated cells are healthy compared to control cells even at the highlight. This data is also supported by the confocal microscopy localization assay for ATG8.

## Conclusion

In this study, we examined the impact of mild osmotic stress (2%PEG) against the highlight stress on *Chlamydomonas reinhardtii* cells. The result of biophysical and biochemical data suggest that low PEG treated cells could combat the highlight stress more effectively than control. Application of such a low concentration of PEG promotes algal biomass without compromising photochemical yield. Our study suggests a novel photoprotective mechanism operating in PEG-treated samples, which has not been reported to date. Since Polyethylene glycol (PEG) is a macromolecule, it cannot penetrate the algal cell wall; also our previous study showed that a higher concentration of PEG inhibited cell growth (reference). Hence we hypothesise that these particles on low concentration might get deposited on the cellular surface, thereby shielding the excess highlight and conferring indirect photoprotection. These findings are promising as algal biomass has immense commercial importance in biofuel, pharmaceutical, cosmetics industries etc.

## Author Contributions

R.S designed the research; JX, R.M.Y performed the research; JX, RMY and R.S analyzed the data; R.S, RMY and JX wrote the paper.

Competing Interest Statement: Authors have no conflicts of interest to declare.

## Acknowledgements

R.S was supported by the Science & Engineering Research Board (CRG/2020/000489), Joint UGC-ISF Research Grant - File No. 6-8/2018 (IC) and Council of Scientific and Industrial Research (No.38 (1504)/21/EMR-ll) Govt. of India, for financial support. DST-FIST and UGC-SAP, Govt. of India, for infrastructure development for the Department of Plant Sciences, University of Hyderabad. SM acknowledged UGC for fellowship (JRF/SRF).

## Figure and Legends

**Table 1:**
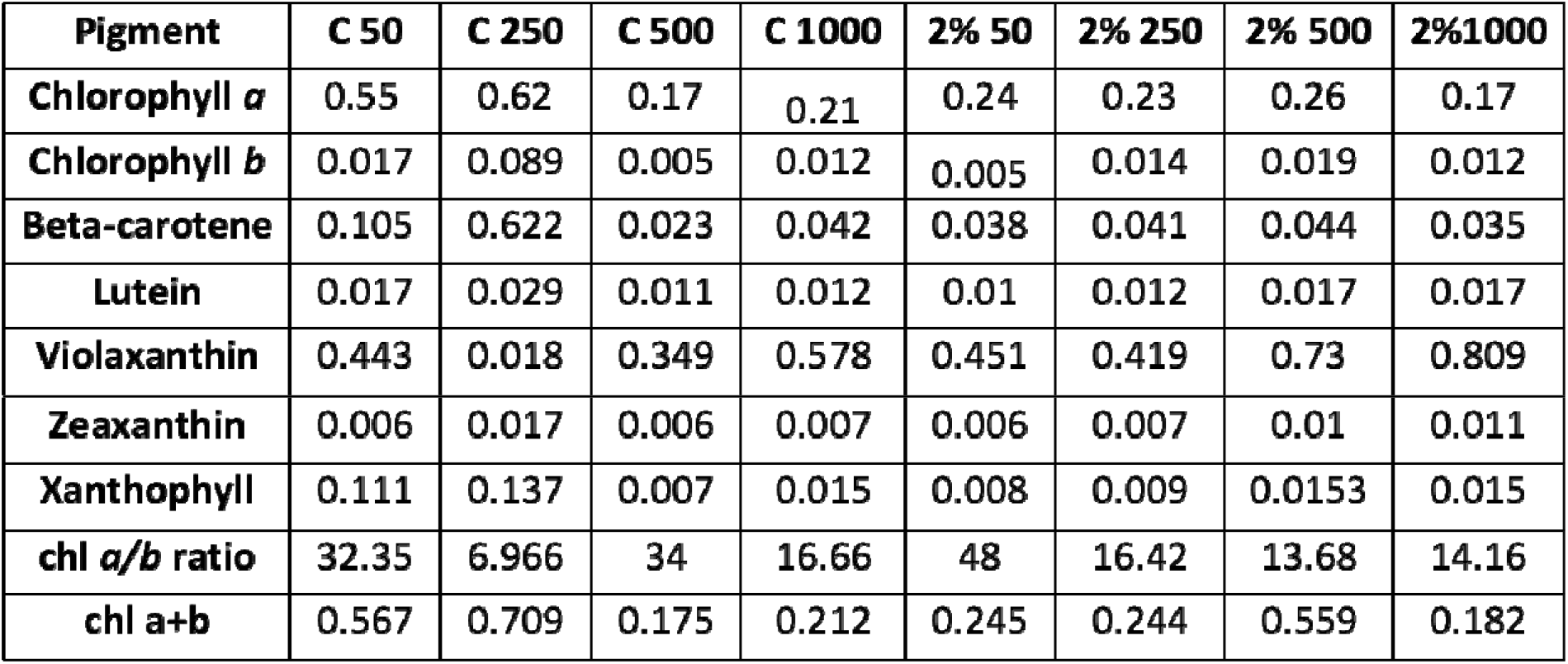
Pigment composition of light and PEG treated samples. Values are mean ± SD (n = 3)

## Supplementary figure

**Fig. S1:**
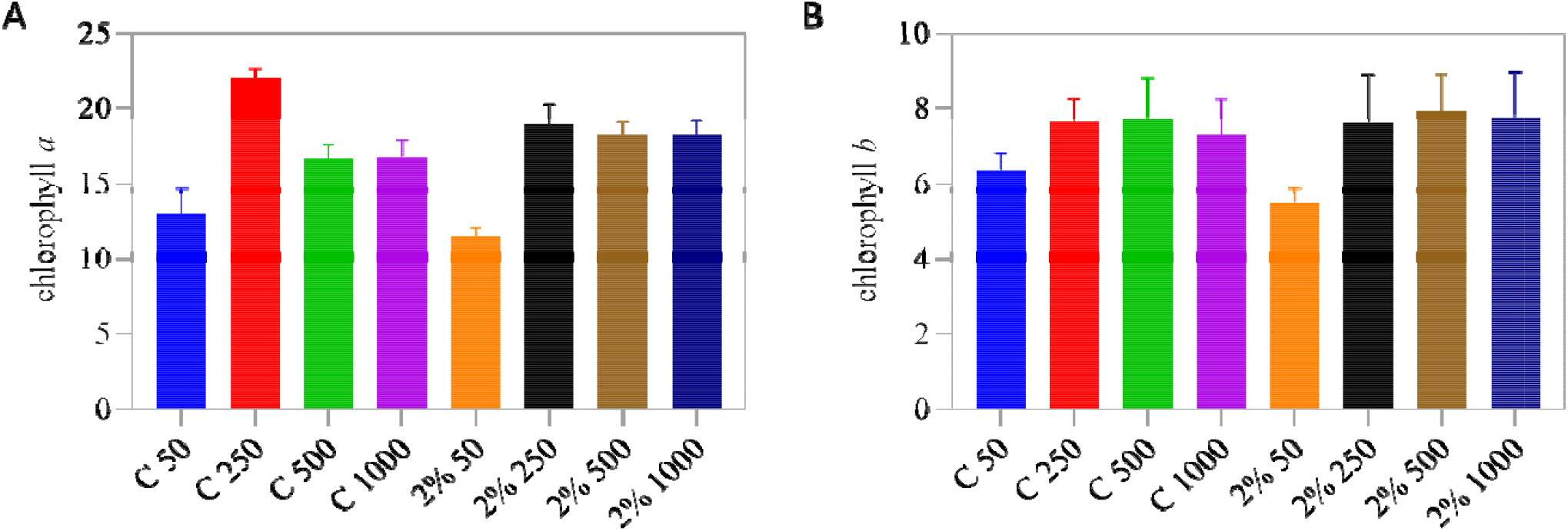
Shows the estimated pigment data calculated by taking triplicates for each samples and readings were taken using PerkinElmer UV visible spectrophotometer and concentration is calculated by adopting equations of lichtenthaler(1987) and Porra et al. (1989). (A) denotes chlorophyll a pigment and (B) denotes chlorophyll b.

**Fig. S2 :**
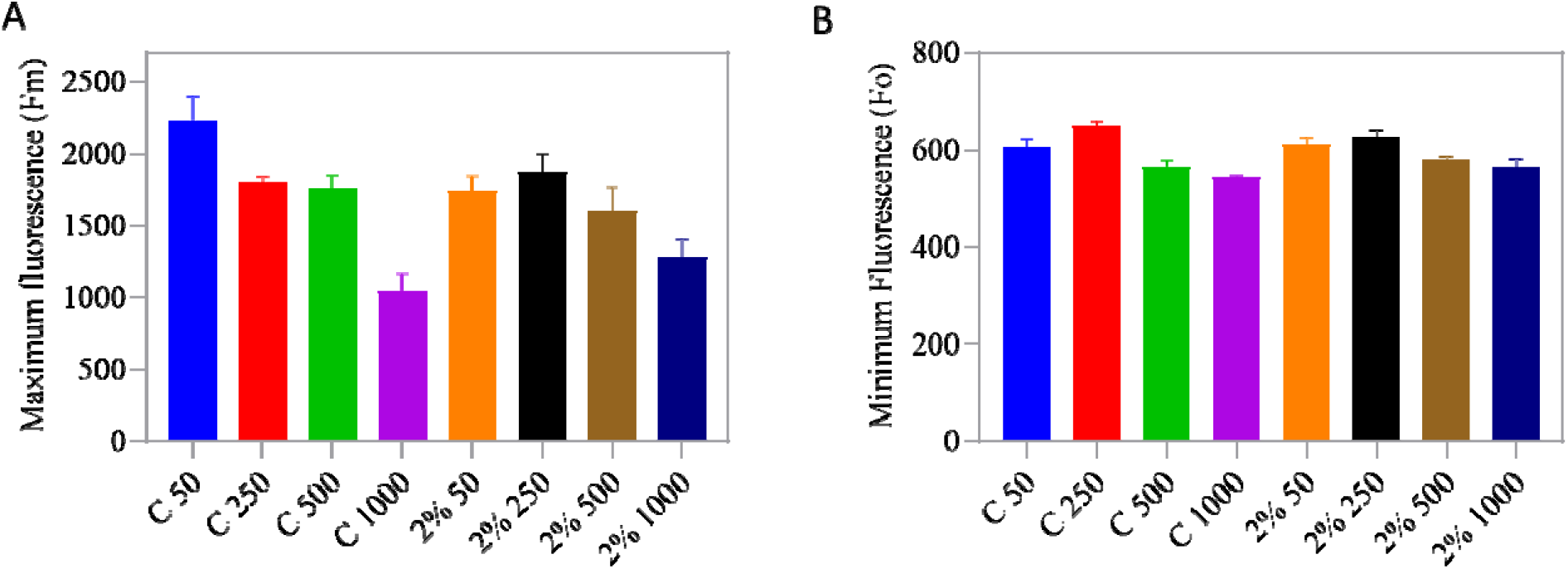
shows the typical OJIP fluorescence transients parameters: fluorescence maximum (Fm) (A) and initial fluorescence (Fo) (B) were calculated by Plant Efficiency Analyzer (Hansatech Instr. Ltd, Kings Lynn, Norfolk, UK) for one second with excitation light wavelength at 650nm in liquid cell cultures.

**Fig. S3:**
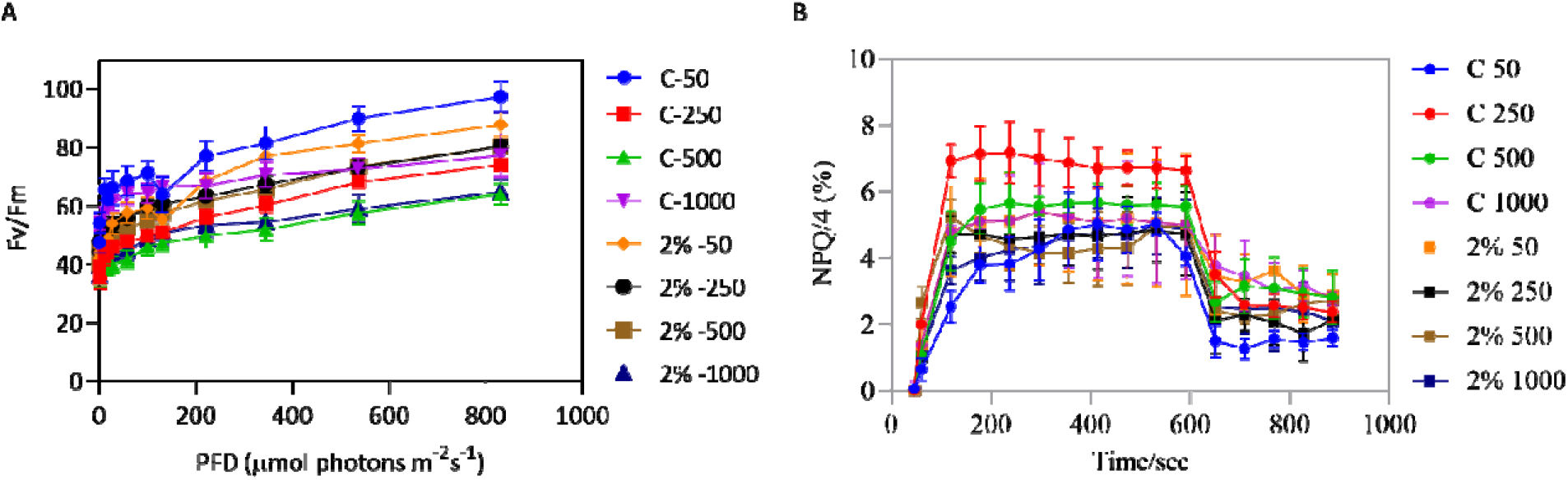
(A) Denotes the photochemical yield in control vs PEG treated samples, (B) gives account of all four NPQ level in each culture condition, calculated by Plant Efficiency Analyzer (Hansatech Instr. Ltd, Kings Lynn, Norfolk, UK) for one second with excitation light wavelength at 650nm in liquid cell cultures.

## Supplementary table

**Table S1:**
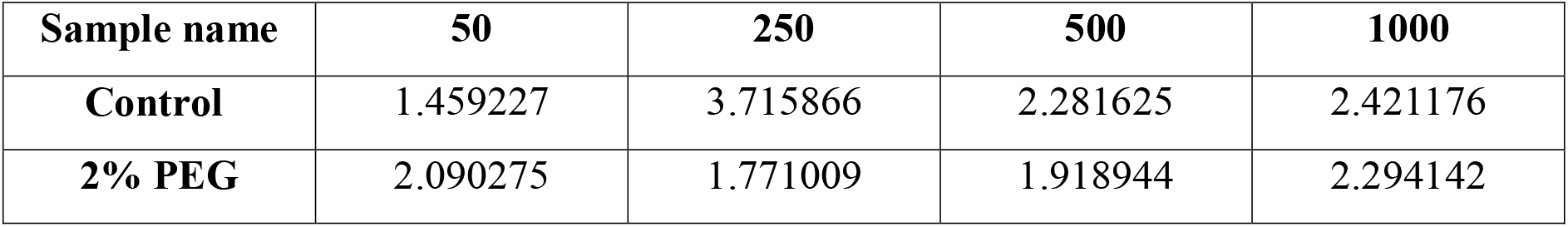
Total carotenoid by OD. represents the carotenoid concentration based on biomass. The samples were taken in triplicates and mean value is considered for all the sample condition.

**Table S2:**
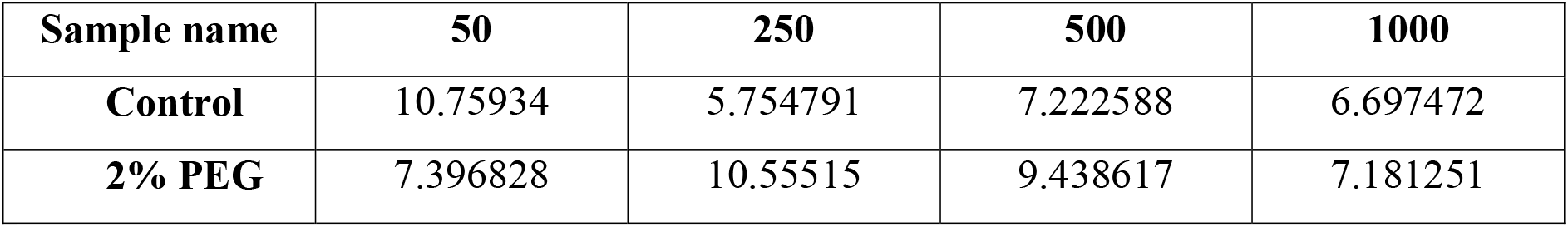
Total chlorophyll and carotenoid. represents chlorophyll vs carotenoid concentration in all culture condition. The samples were taken in triplicates and mean value is considered for all the sample condition.

| **Sample name** | **50** | **250** | **500** | **1000** |
| --- | --- | --- | --- | --- |
| **Control** | 10.75934 | 5.754791 | 7.222588 | 6.697472 |
| **2% PEG** | 7.396828 | 10.55515 | 9.438617 | 7.181251 |

**Table S3:** Total chlorophyll by OD. represents the total chlorophyll concentration based on biomass. The samples were taken in triplicates and mean value is considered for all the sample condition.

| **Sample name** | **50** | **250** | **500** | **1000** |
| --- | --- | --- | --- | --- |
| **Control** | 15.70032 | 21.38403 | 16.47924 | 16.21576 |
| **2% PEG** | 15.4614 | 18.69327 | 18.11218 | 16.47481 |

## Materials and methods

### Growth and experimental culture condition

The *Chlamydomonas reinhardtii* wild-type strain CC125 received from Chlamy center, USA, were grown photoheterotrophically in Tris-acetate phosphate medium(TAP) under controlled conditions in the laboratory with continuous illumination of the white light of 50 μmol photons m^−2^s^−1^ at 25^0^ C temperature in Algae Tron growth chamber, Czech Republic. When the mother culture reaches an optical density of around 1, it is inoculated into the control medium without PEG and the treatment medium having 2%PEG. Both the sets will be grown at different intensities of light (50, 250, 500 and 1000 μmol photons m^−2^s^−1^).

#### Growth curve and ROS Measurement

The cell density of all samples was examined at intervals of 24, 48 and 72 hrs using a UV-visible spectrophotometer by measuring OD at 750nm; the TAP medium is kept blank. It was reported that at 750nm, the wavelength was not absorbed by chlorophyll and carotenoid pigments hence reading reflects only light scattering by cells which is proportional to cell density(Yadav et al., 2020). The 2,7-dichlorodihydrofluorescein diacetate (H_2_DCFDA) (Sigma-Aldrich), a fluorescence dye, is used to detect total ROS in live *C. reinhardtii* cells. Cells at a 3 million density were harvested from all the conditions, and H_2_DCFDA staining was performed at 10 μM concentration (Upadhyaya et al., 2020). Further, cells were incubated with dye for 1 h at RT in a continuously rotating shaker under dark. Images were captured using Carl Zeiss NL0 710 Confocal microscope. H_2_DCFDA was detected in a 500–530 nm bandpass optical filter with an excitation wavelength of 492 nm and an emission wavelength of 525 nm. Chlorophyll auto-fluorescence was detected using an optical filter of 600 nm. Samples were viewed with a 60× oil immersion lens objective by using the ZEN 2010 software.

#### Chlorophyll and HPLC Pigment Analysis

Each culture was centrifuged at 5000 rpm for 5 minutes, and pellets were treated with 1 mL of 80% acetone. Vortex the mixture and incubate it at -20^0^ C for 1hr. Pellet down them at 10000rpm and take OD values at 480nm,510nm(carotenoid) and 645 nm, 663 nm(chlorophyll) using a UV-visible spectrophotometer (Perkin Elmer). Total chlorophyll, chl *a*, chl *b* and carotenoid ratio is calculated at 24, 48 and 72 hrs intervals based on the equations of (Benitez, 1989; Lichtenthaler, 1987; Porra et al., 2019). The samples were harvested around 0.6-0.8 OD at 750nm and will be incubated with 100% acetone overnight for pigment extraction. These samples were centrifuged at 10,000 rpm for 10 minutes, and the supernatant was filtered by a 0.45μm filter. Shimadzu HPLC will be used to separate the pigments on the C-18 column (250 × 4.6 mm, 5μm; Phenomenex). The isocratic system of methanol: acetonitrile: acetone (70:20:10) will be used as the mobile phase. Each pigment will be estimated quantitatively using the respective standard.

#### Fast chlorophyll, a fluorescence measurement

Plant Efficiency Analyzer (Hansatech Instr. Ltd, Kings Lynn, Norfolk, UK) was used to measure OJIP fluorescence transients of all samples under study for one second with an excitation light wavelength of 650 nm in liquid cell cultures. The maximum quantum yield of PSII (Fv/Fm) was calculated by the Handy PEA instrument at 3000 μmol photons m^−2^ s^−1^ light intensity using the following formula Fv/Fm = (Fm – Fo)/Fm, where Fv is the variable fluorescence, Fo is the initial fluorescence level recorded at 50μs after the onset of illumination, and Fm is the maximum fluorescence. Minimum fluorescence (Fo) (i.e. measured in the dark-adapted state) ([PDF] The fluorescence transient as a tool to characterize and screen photosynthetic samples | Semantic Scholar, n.d.). The electron transport efficiency between Q_A_ and Q_B_ quinones of PSII can also be determined by the using parameter like 1 − VJ = 1-(FJ − F0)/ (Fm − F0), where FJ is the fluorescence level at 2 ms after the onset of illumination (Kodru et al., 2015).

#### Measurements of chlorophyll fluorescence and light curve-based parameter

The Chl fluorescence and light curve-based parameter were determined by Dual-PAM 100 (Walz, Germany) in control and PEG-treated samples. Each culture was dark-adapted for 20 min before the PAM parameter analysis. To perform quantum yield measurements, we recorded a light response curve by changing the actinic light intensity (stepwise PAR ranges from 3 µmol photons m ^−2^ s ^−1^ - 1500 µmol photons m ^−2^ s ^−1^). While recording light response curves, samples were exposed to each light intensity for < 4 min followed by a saturating pulse (SP) of 4000 µmol photons m ^−2^ s ^−1^. Electron flow through PSII was calculated as ETR(II) = 0.84 × 0.5 × Y(II) × light intensity (μmol photons m ^−2^ s ^−1^) (Krall and Edwards, 1992). The quantum yield of PSII; Y(II), was calculated as (Fm’ – Fs) / Fm’. Fs denotes the steady-state fluorescence, while Fm and Fm’ are the maximum fluorescence levels in the dark and the light, respectively. Fo’ was calculated as Fo / (Fv / Fm + Fo / Fm’) (Oxborough and Baker, 1997). The fraction of non-regulatory quantum yield of energy dissipation is calculated as Y(NO) = Fs / Fm; the quantum yield of regulated energy dissipation of PSII, Y(NPQ) = 1-Y(II) - Y(NO). The chl fluorescence for NPQ measurements, control and PEG-supplemented cells were grown in a TAP medium for 72 h. After dark adaptation, cells were pre-illuminated for 2 min with a weak (3 µmol photons m^−2^ s^−1^) far-red LED; maximum fluorescence (F_m_) and changes in maximal fluorescence in light (F_m_‵) are measured by applying saturating pulse for 10 min induction and 5 min relaxation. NPQ was calculated using the formula NPQ = (F_m_-F_m_‵)/ F_m_‵; actinic light was 660 µmol photons m^−2^ s^−1^ and saturating light, 8000 µmol photons m^−2^ s^−1^. The far-red LED was kept on during dark recovery to oxidize PSI and prevent over-reduction of the PQ pool (Bonente et al., 2011).

#### Localization of ATG8 using Immunofluorescence Microscopy

The 3 × 10^6^ cells/ml of cells were fixed in 4% paraformaldehyde and 15% sucrose dissolved in phosphate saline buffer (PBS) for 1 h at RT, later, the cells were washed twice with PBS buffer. Further cells were permeabilized by incubation with 0.01% Triton X100 in PBS for 5 min at RT and washed twice with PBS. Next, the samples were transferred to sterile Eppendorf tubes and blocked with a 1% BSA (w/v) in PBS for 1 h. These Samples were incubated with anti-ATG8 primary antibody diluted (1:1000) in PBS buffer, pH = 7.2, containing 1% BSA overnight at 4°C on a rotatory shaker. Cells were then washed twice with PBS for 10 min at 25°C, followed by incubation in a 1:10000 dilution of the fluorescein Dylight 405 labelled goat anti-rabbit secondary antibody (Sigma) in PBS-BSA for 2 h at 25°C. These cells were washed three times with PBS for 5 min, and Images were captured with Carl Zeiss NL0 710 Confocal microscope. ATG8 with the following parameters: excitation of Dylight 405 nm and emission at 420 nm; images were analyzed with ZEN software(Chouhan et al., 2022).

#### Immunoblotting for Thylakoid membrane protein

The global proteome of each sample is separated by SDS-PAGE loaded with equal chlorophyll concentration. Separated proteins were transferred to polyvinylidene difluoride (PVDF) membrane from Bio-Rad company using transblot apparatus (Bio-Rad). Primary antibodies against LHCII, PSII, and PSI complex proteins were Lhcb5purchased from Agrisera. The primary antibody dilutions are as follows: D1, Lhcb4, PsaG, PsaH, Lhca1, Lhca2 and LhcSR3(**1:10000)**, D2, PsaA, PsaF, ATG8, Lhcb1, Lhcb5 and Lhcb2 (**1:5000**). Followed by incubation with secondary antibody ligated to horse radish peroxidase (**1:10000**) dilution. The images were developed using Bio-Rad’s New Chemi Doc™ Touch Imaging System (Nama et al., 2018).

### Circular Dichroism

CD spectra were recorded using JASCO 815 spectropolarimeter within the 400-750 nm wavelength range and a scan speed of 100 nm min^−1^ with 3 cumulative accumulations. Optical pathlength 1 cm, bandwidth 2nm and a data pitch of 0.5 nm were used.

### Statistical analysis

The measurements were carried out on randomly selected *C. reinhardtii* samples, and the data collected are the mean ± SD of a minimum of 3 and a maximum of 5 biological replicates. All graphs were plotted using Prism 8.0 software.

